# Novel mediation analysis of human plasma proteome and metabolome reveals mediators of improved glycemia after gastric bypass surgery

**DOI:** 10.1101/817494

**Authors:** Jonathan M Dreyfuss, Yixing Yuchi, Hui Pan, Xuehong Dong, Donald C. Simonson, Ashley Vernon, Pratik Aryal, Anish Konkar, Yinong Sebastian, Brandon W Higgs, Joseph Grimsby, Cristina M. Rondinone, Simon Kasif, Barbara B. Kahn, Kathleen Foster, Allison Goldfine, Mary-Elizabeth Patti

## Abstract

Molecular mechanisms by which Roux-en-Y gastric bypass (RYGB) improves glycemic control and metabolism in type 2 diabetes (T2D) remain incompletely understood. In the SLIMM-T2D trial, participants with T2D were randomized to RYGB or nonsurgical management and their fasting plasma proteome and metabolome were analyzed for up to 3 years. To identify analytes that mediate improvement in outcomes, we developed a high-throughput mediation analysis method (Hitman), which is significantly more powerful than existing methods. Top-ranking analyte mediators of glycemia improvement were growth hormone receptor and prolylhydroxyproline, which were more significant than any clinical mediator, including BMI. Beta-alanine and Histidine Metabolism (both including CNDP1) were top differentially regulated pathways, and Valine, Leucine and Isoleucine Degradation was also a top differentially-regulated pathway and a top mediator of improvement in insulin resistance. The identified analytes may serve as novel targets for T2D therapy. More broadly, Hitman can identify analyte mediators of outcomes in randomized trials for which high-throughput data are available.

## Introduction

Bariatric surgery is a potent approach to manage obesity and related comorbidities, including type 2 diabetes (T2D) and its complications (*1–3*). Roux-en-Y gastric bypass (RYGB) has particularly powerful effects on glucose metabolism, resulting in remission of T2D in about 90% of patients at 1 year and 45% at 5 years (*4–10*). Improved glycemic control occurs days after RYGB, before substantial weight loss, supporting that weight loss-independent mechanisms drive metabolic improvements (*11–13*). Moreover, recent observational studies indicate that bariatric surgery is associated with fewer microvascular and macrovascular complications of diabetes and reduced mortality (*14–16*). Identifying molecular mechanisms responsible for improved systemic metabolism could allow development of novel nonsurgical approaches for T2D treatment.

Improved postoperative glycemic control is linked to enhanced meal-related insulin secretion and increases in incretin hormones such as GLP-1 and PYY (*17–19*). However, preclinical studies indicate incretins are not required for beneficial effects of bariatric procedures (*20–24*). Other postoperative changes include reductions in branched chain amino acids (*BCAA*)(*25–27*) and aromatic amino acids (*28*), alterations in plasma bile acids and FXR signaling (*29–31*) and in other hormones (*32*). RYGB alters the microbiome community composition toward that of less obese individuals (*33*), with increased abundance of many amino acid fermenters (*34*). Yet, microbiome changes cannot fully explain clinical improvement (*35*). Thus, the primary molecular factor(s) contributing to improved metabolism and diabetes remission remain uncertain.

We assayed fasting plasma samples from the Surgery or Lifestyle with Intensive Medical Management in the Treatment of Type 2 Diabetes (SLIMM-T2D) clinical trial (clinicaltrials.gov:NCT01073020), in which 38 obese participants with T2D were randomized to RYGB (n = 19) or nonsurgical intensive diabetes weight management (DWM; n = 19) and followed longitudinally for 3 years (*10, 36*). Groups were similar at baseline (pre-randomization). Greater clinical improvement was seen in RYGB than in DWM for multiple outcomes, including body weight reduction (assessed by BMI), glycemia (assessed by HbA1c), triglycerides, HDL cholesterol, systolic blood pressure, and reductions in the number of antidiabetic, antihypertensive, and lipid-lowering medications (Simonson et al., 2018). All RYGB participants achieved 10% weight loss before 3 months, whereas 37% of DWM participants achieved 10% weight loss before 3 months (*36*). To identify candidate molecules responsible for improvement in glucose metabolism after RYGB as compared with DWM, we developed novel tools and applied them to clinical, proteomic, and metabolomic data from these samples.

## Results

### Clinical

Proteomic and metabolomic data at baseline, the 3 month time point, and 1 year were available from 35 of the 38 SLIMM-T2D participants. Metabolic characteristics of these 35 participants did not differ between groups at baseline (**Table S1**). The HbA1c and BMI of those that had proteomics or metabolomics per time point also did not differ from the SLIMM-T2D participants within their group at any time point (**Figure S1)**. After 3 years, no DWM participants achieved study-defined glycemic goals (HbA1c<6.5% and fasting plasma glucose<126 mg/dL), whereas eight RYGB participants did, and seven of these eight participants were not receiving any antidiabetes medications (Simonson et al., 2018). The number of participants taking each diabetes medication class per arm at each time point is tabulated in **Table S1**.

### Proteome

Fasting proteomics were profiled at baseline, the 3 month time point, and years 1, 2 and 3 in RYGB and DWM. At baseline (pre-randomization), there were no statistically significant proteins as assessed by false discovery rate (FDR<0.15). For later, post-randomization time points, reduction in BMI was greater in RYGB, so groups were compared both with and without adjustment for each person’s BMI change. Differences in the baseline-corrected proteome in RYGB vs. DWM (i.e. differences between groups in changes over time) emerged at the 3 month time point, with 14 significant proteins in the unadjusted analysis and 8 in the BMI-adjusted analysis. Proteins common to both analyses and downregulated in RYGB were: carnosine dipeptidase 1 (CNDP1, also known as Beta-Ala-His dipeptidase 1) and Fetuin-B (FETUB). Proteins in common upregulated in RYGB were: integrin-binding sialoprotein (IBSP), insulin-like growth factor binding protein (IGFBP2), endothelial cell-specific molecule 1 (ESM1), macrophage metalloelastase (MMP12), alpha-1-antichymotrypsin complex (SERPINA3), and CC motif chemokine 22 (CCL22).

Differences of baseline-corrected protein abundance in RYGB vs. DWM persisted at years 1, 2, and 3. The log_2_ fold change values in RYGB vs. DWM for the 19 proteins differentially abundant with fold-change magnitude at least 1.5 after baseline correction at any time point without BMI adjustment are depicted in the heatmap in **Figure 1**. Statistics for all comparisons in all analytes are presented in **Table S2.** Upregulation of IGFBP2 (>50%, p<0.001 at all time points in SOMAscan, **Figure S2A**) was confirmed by ELISA, which showed significant differences in baseline-corrected values (p<0.05) at 12, 24, and 36 months (**Figure S2B**), and ELISA changes over time were significantly correlated to corresponding SOMAscan changes (r=0.78, p<10^-7^).

**Figure 1.**
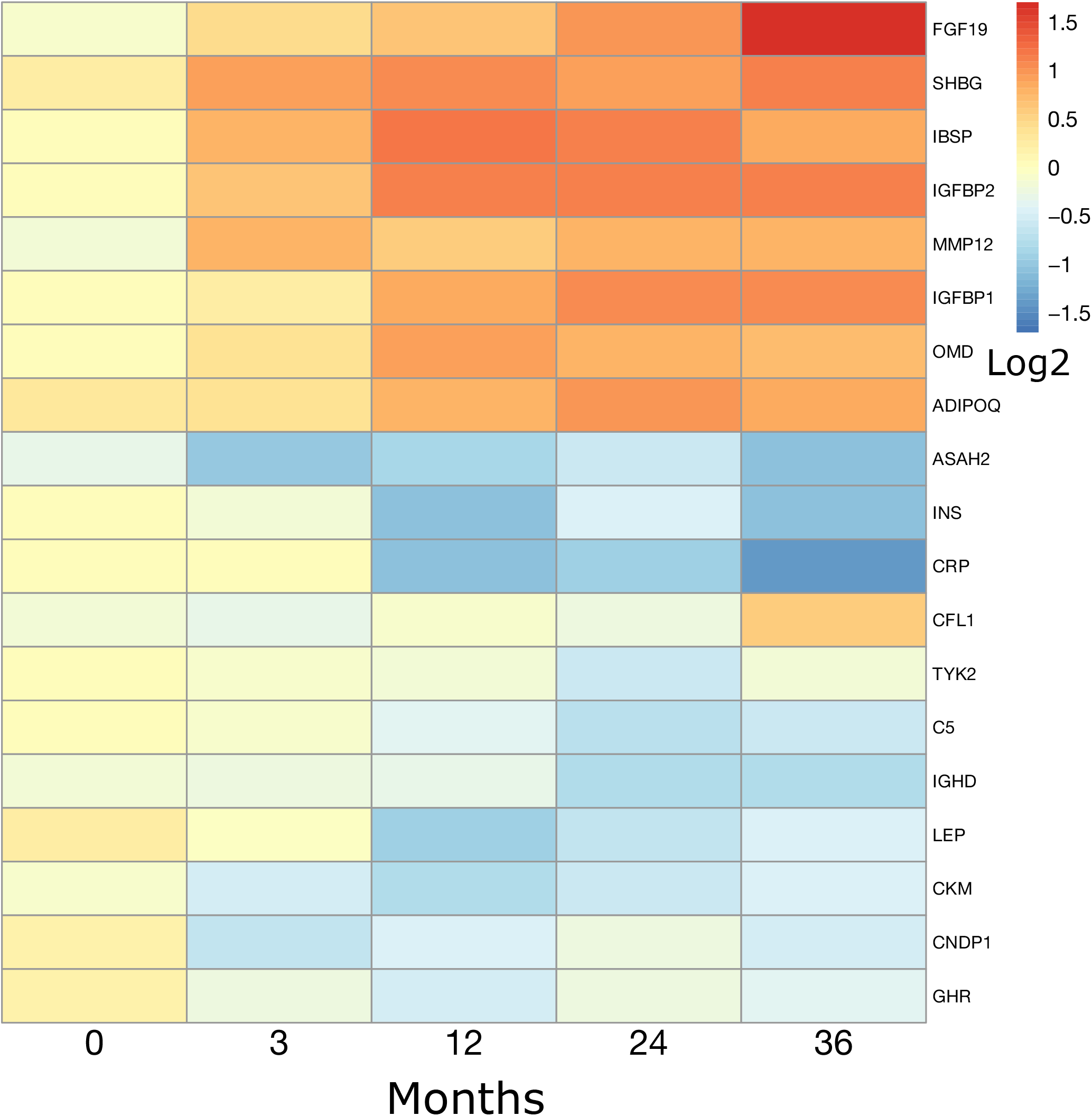
Proteome changes, see also Figure S2 and Table S2. Heatmap of log_2_(RYGB/DWM) at all time points (post-baseline log_2_ abundance values are baseline-corrected) for proteins that are differentially abundant at any time point (FDR<0.15 and foldchange magnitude > 1.5).

### Metabolome

At baseline (pre-randomization), there were no significant differences in the fasting metabolome between groups. Significant differences in the baseline-corrected metabolome in RYGB vs. DWM emerged at the 3 month time point, with 96 metabolites changed in the unadjusted analysis and 74 in the BMI-adjusted analysis; 45 of the metabolites were common to both analyses.

Differences of baseline-corrected metabolite abundance in RYGB vs. DWM persisted at years 1, 2, and 3. The log_2_ fold change values in RYGB vs. DWM for the 85 metabolites differentially abundant with fold-change magnitude at least 1.5 after baseline correction at any time point without BMI adjustment are depicted in the heatmap in **Figure 2**. The color range was defined to match **Figure 1**, with fold-change magnitudes above 3 (|log_2_ fold change| > 1.6) treated as fold-change magnitude of 3, the limit of the color bar. However, we saw much larger foldchanges in the metabolome. For example, the BCAA-related metabolite 3-hydroxyisobutyrate was 88% lower (i.e. down by >9-fold) at the 3 month time point in RYGB than DWM, and it was also significant in the BMI-adjusted analysis. Statistics for all comparisons in all analytes are presented in **Table S2.** For the metabolites in **Figure 2**, we tested the correlation of their change from baseline at the 3 month time point to the corresponding changes in the proteins shown in **Figure 1** to identify co-regulation. These correlations are provided in **Table S2**.

**Figure 2.**
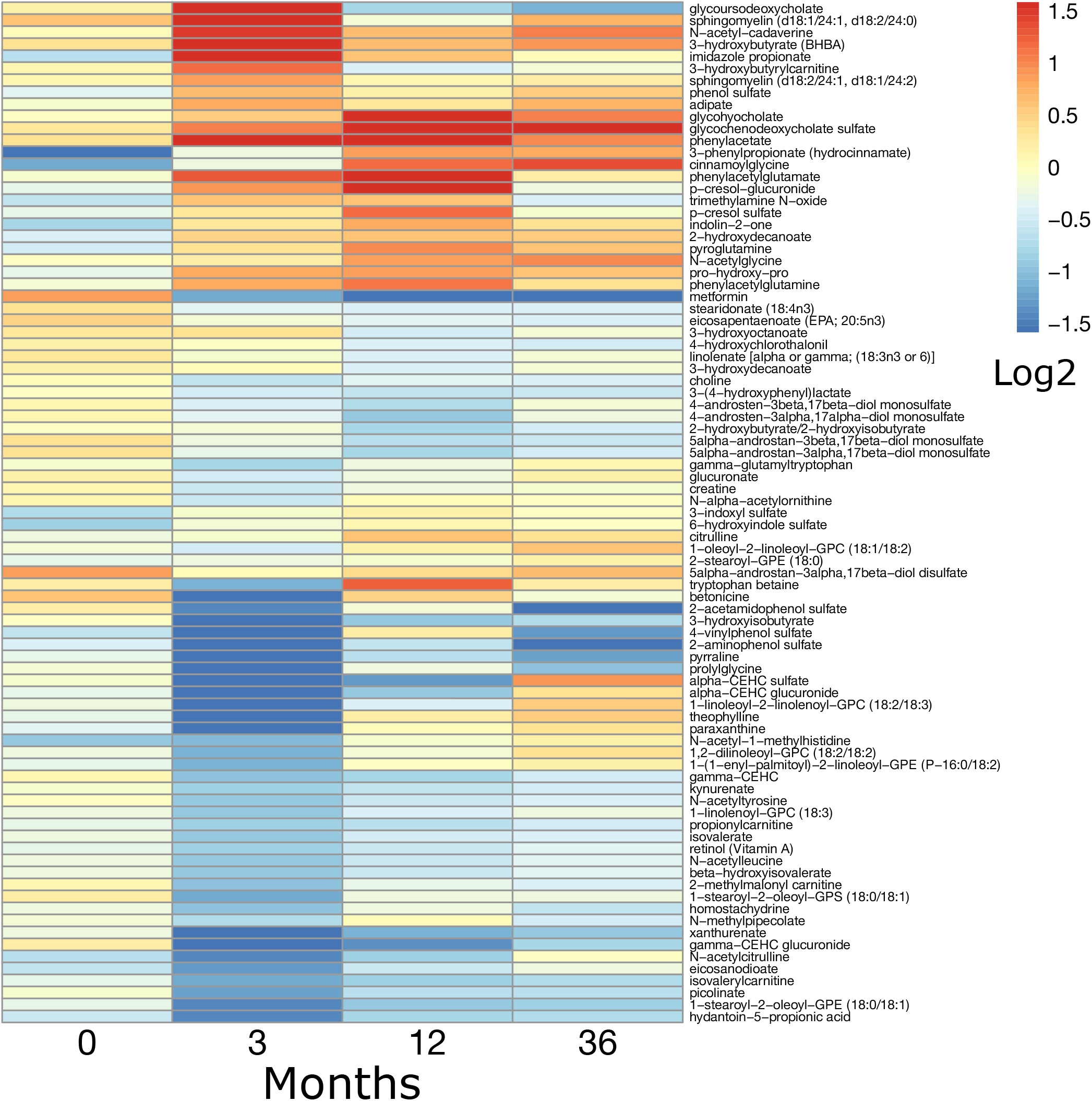
Metabolome changes, see also Table S2. Heatmap of log_2_(RYGB/DWM) at all time points (post-baseline log_2_ abundance values are baseline-corrected) for metabolites that are differentially abundant at any time point (FDR<0.15 and fold-change magnitude > 1.5). Log_2_ values outside the range −1.58 to 1.58 are shrunken toward zero to be in this range.

### Proteomic and metabolomic integrative pathway analysis

We integrated proteomics and metabolomics for pathway analysis by creating a single, integrated dataset, and testing pathways composed of both proteins and metabolites from the Small Molecule Pathway Database (*37*). We identified differentially abundant pathways between groups without BMI adjustment at the 3 month time point. The 3 month time point showed robust improvements in glucose metabolism in both groups, yet preceded larger weight differences. There were 50 significant pathways.

Top-ranking pathways are presented in **Figure 3A**. The top-ranking pathway is Phospholipid Biosynthesis (**Figure 3B**), whose top analytes were choline (**Figure S2C**) and choline phosphate (**Figure S2D**), both reduced in RYGB relative to DWM. The second pathway was Valine, Leucine, and Isoleucine Degradation, whose top analytes were the valine catabolic intermediate 3-hydroxyisobutyrate acid (88% lower at the 3 month time point in RYGB; **Figure 4**, middle row, right), the ketoacid 3-methyl-2-oxobutyrate (ketoisovalerate; **Figure 4**, middle row, second from right), and the BCAAs valine and leucine (**Figure 4**, top row), all with sustained reductions after RYGB. Similar reductions were observed for other ketoacids and downstream acylcarnitines (**Figure 4**). Propionylcarnitine, a C3 acylcarnitine product of valine and isoleucine metabolism (**Figure 4**, lower row) with a 50% reduction in RYGB, was the top analyte of the related pathway Oxidation of Branched Chain Fatty Acids.

**Figure 3:**
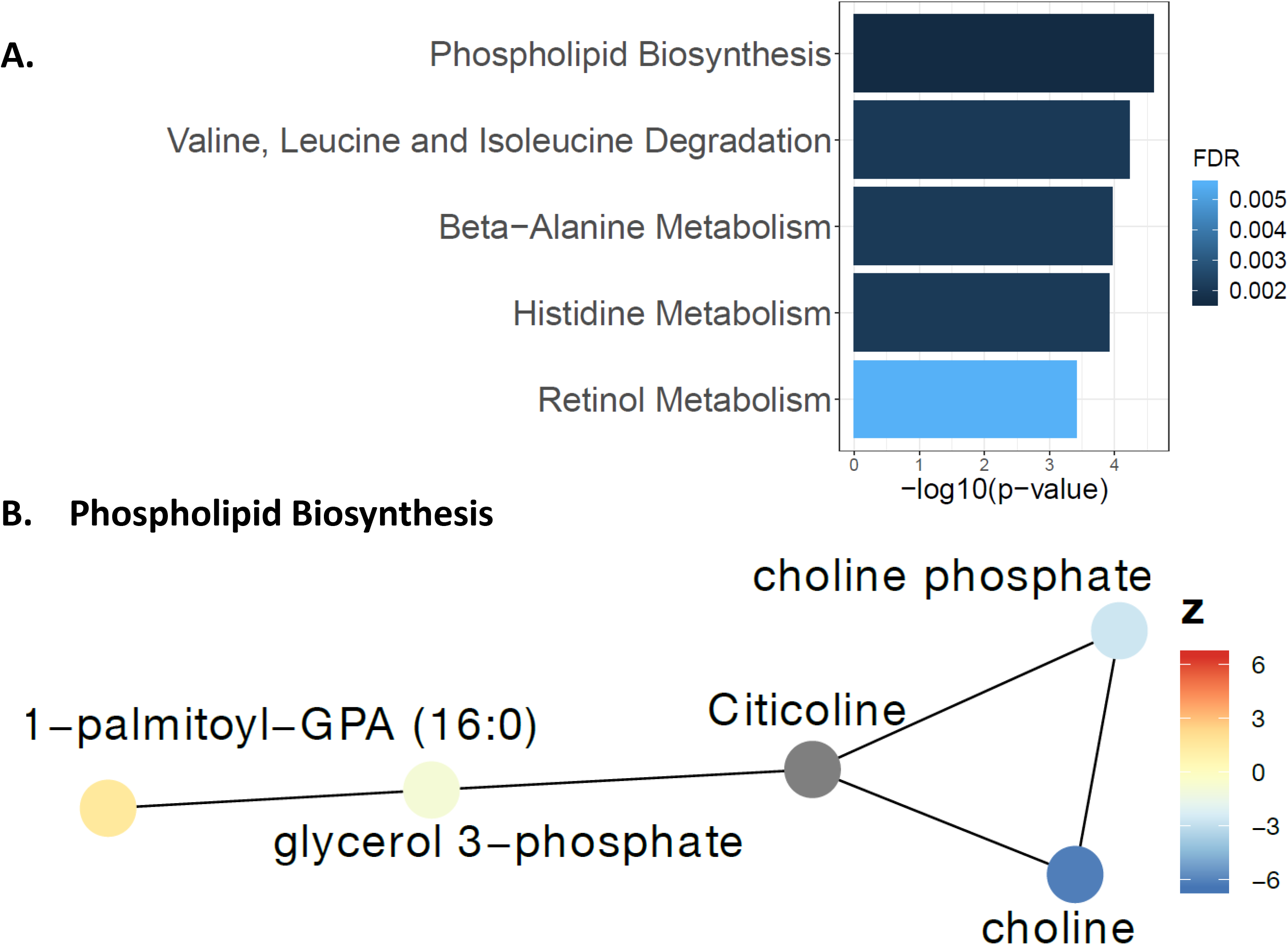
Top differential pathways and Phospholipid Biosynthesis network. (A) –log_10_(p-values) and false discovery rates (FDRs) of top pathways. (B) Nodes are colored by between-group z-score, whereas unmeasured nodes are colored dark gray. Connections are from Pathway Commons network.

**Figure 4.**
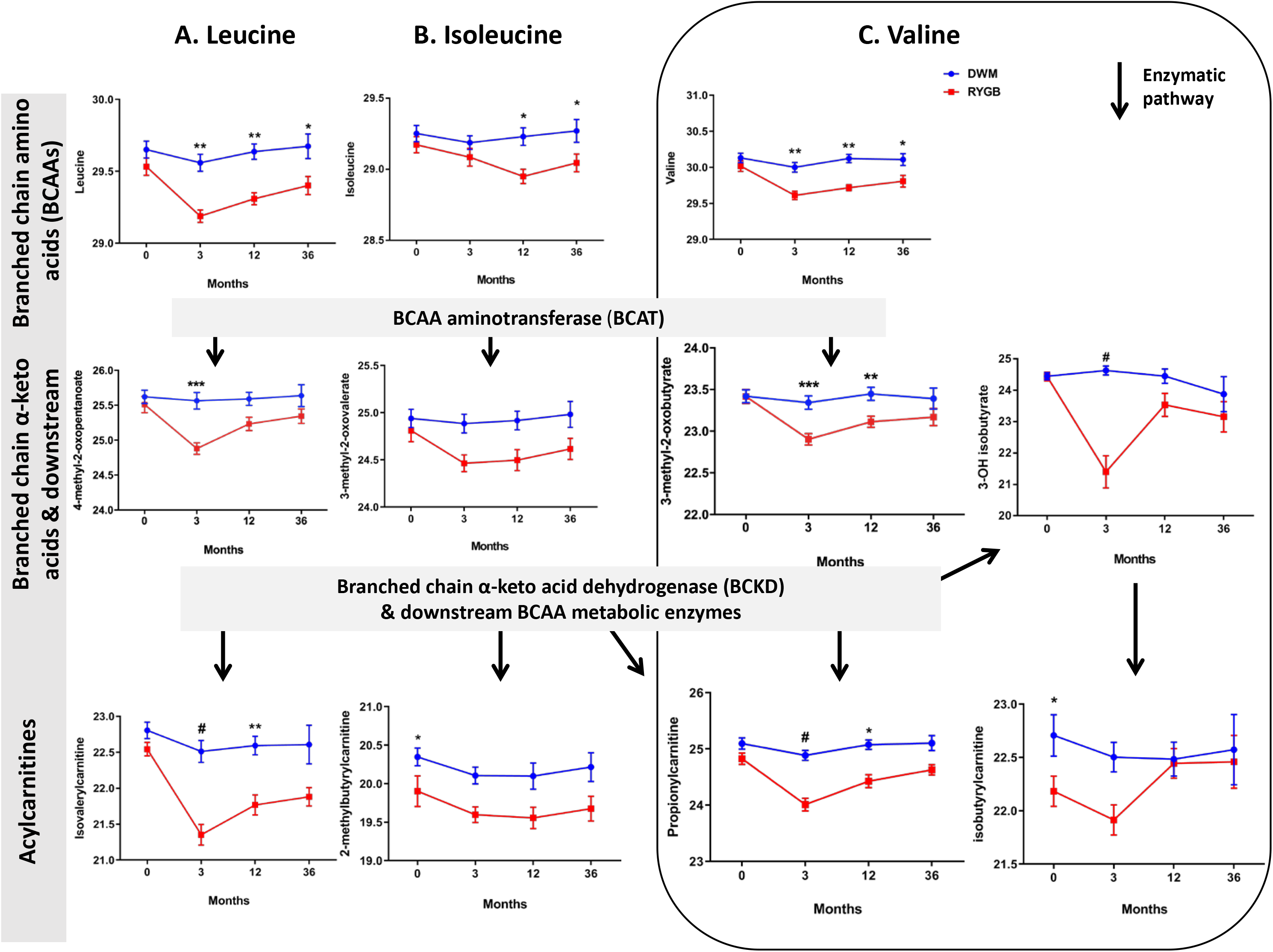
Branched chain amino acids (BCAA) and downstream metabolites, see also Table S2. Arrows indicate enzymatic pathway relationships. Data were analyzed by moderated t-tests; post-baseline time points were analyzed using change from baseline. Data are reported as mean ± SEM on the log2 scale. *p<0.05, **p<0.01, ***p<0.001, and #p<0.0001.

Beta-alanine Metabolism’s and Histidine Metabolism’s (**Figure 5A**) top analytes were CNDP1 and its enzymatic product histidine. Histidine was reduced after RYGB but not DWM (FDR 0.002, **Figure 5D**), and its reduction was correlated with the reduction of CNDP1 (r=0.40, p=0.02). CNDP1 was reduced by 43% after RYGB at the 3 month time point (FDR=1.6*10^-5^) and remained lower in RYGB at 36 months (**Figure 5B**). A previous study had been unable to validate SOMAscan CNDP1 measurements (*38*). ELISA-determined CNDP1 levels in our analysis showed little correlation with SOMAscan levels per time point, but changes within person over time showed stronger correlation (r=0.21, p=0.055). Moreover, we confirmed significant reductions in CNDP1 after RYGB, demonstrating a 68% decrease in baseline-corrected change for RYGB vs. DWM (p=0.004 at the 3 month time point, **Figure 5C**).

**Figure 5.**
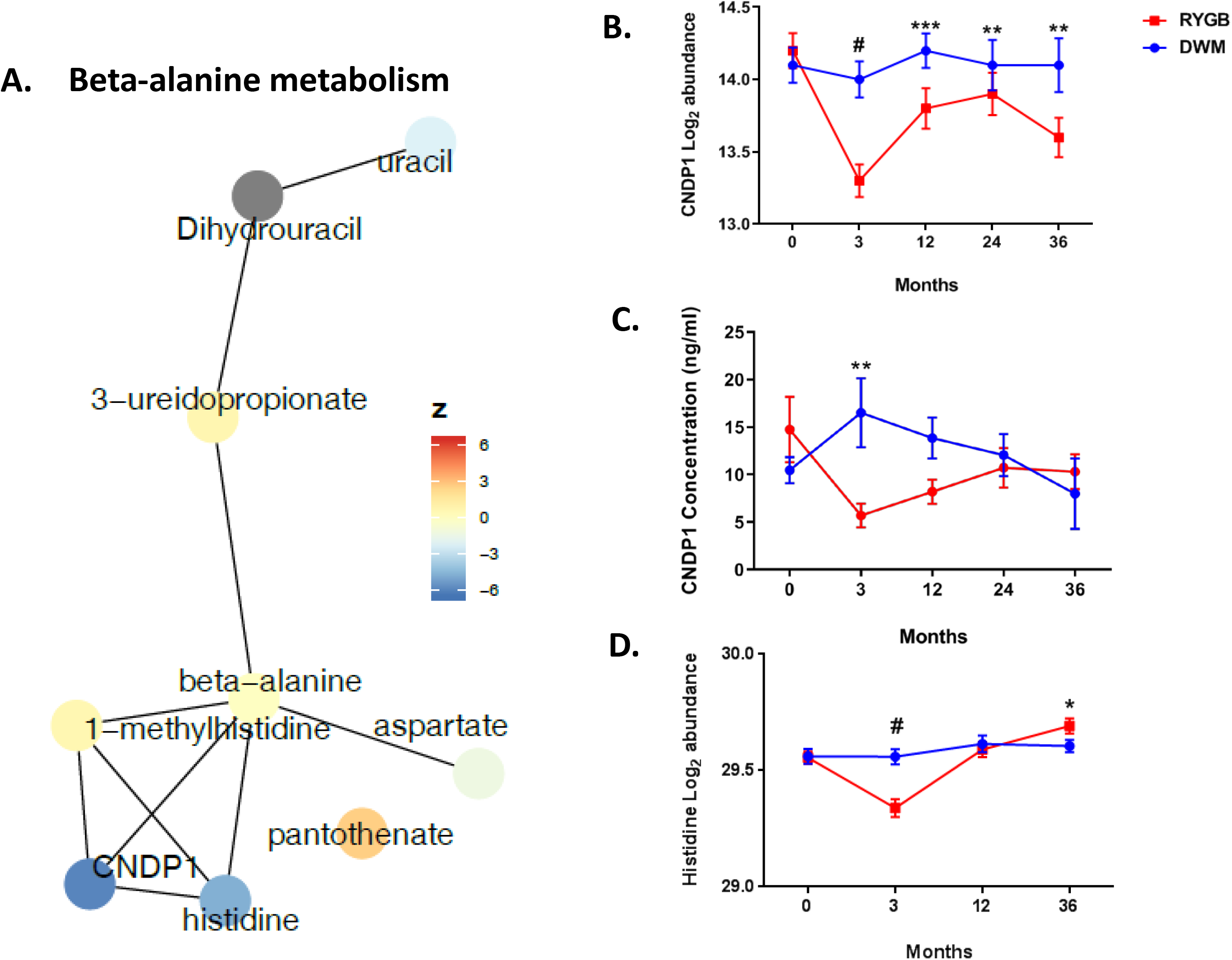
Beta-alanine metabolism, see also Table S2. (A) Network where nodes are colored by between-group z-score, whereas unmeasured nodes are colored gray. Connections are from Pathway Commons network. (B) Log_2_ abundance of CNDP1 measured by SOMAscan. (C) CNDP1 plasma levels measured by ELISA. (D) Log_2_ abundance of histidine measured by metabolomics. Data were analyzed by moderated t-tests; post-baseline time points were analyzed using change from baseline. Data in B, C, and D are reported as mean ± SEM. *p<0.05, **p<0.01, ***p<0.001, and #p<0.0001.

Retinol Metabolism’s top analyte was retinol (vitamin A), whose baseline-corrected abundance decreased by 33% in RYGB vs. DWM at the 3 month time point (FDR<10^-4^), and remained nominally lower at 12 months (p=0.02, FDR=0.18), but reverted to baseline values in both groups at 36 months (Figure S3A). Retinol binding protein 4 (RBP4) tended to be decreased in RYGB (10% lower at the 3 month time point, **Figure S3B**); these changes in RBP4 were significantly correlated to changes in retinol over all time points (r=0.33, p=0.001). Given the role of RBP4 as a mediator of insulin resistance (*39*), we measured RBP4 and its partner transthyretin (TTR) in a random subset of 12 subjects by quantitative western blot at multiple time points. At the 3 month time point, RBP4 and TTR were numerically lower in RYGB vs. DWM, but differences did not reach statistical significance with this sample size (**Figure S3C, S3D**). However, RBP4 changes by western blot correlated significantly to changes by SOMAscan (r=0.28, p=0.03), and changes in both RBP4 and TTR significantly correlated to each other (r=0.63, p<10^-5^) and to changes in retinol (both r>0.6, p<10^-4^).

### High-throughput Mediation Analysis (Hitman)

We sought to identify analytes and clinical measures at the 3 month time point that mediated RYGB’s improvement in glycemia at 1 year. While there are numerous mediation methods, two previous comprehensive comparisons, including one for genome-wide data, concluded that the best balance between false positive rate and power was offered by the joint significance method (*40, 41*). This method was mathematically shown to control its false positive rate and to be more powerful than some of its peers (*42*). A drawback of the joint significance test is that it does not account for direction of effect of the mediator, so it could call as significant an “inconsistent” mediator (*43*). For example, RYGB (compared to DWM) decreases retinol at the 3 month time point and HbA1c at 1 year. However, a decrease in retinol is associated with an *increase* in HbA1c (**Figure 6, left**). Thus, the direction of retinol’s mediation is inconsistent with RYGB-mediated improvement in HbA1c.

**Figure 6.**
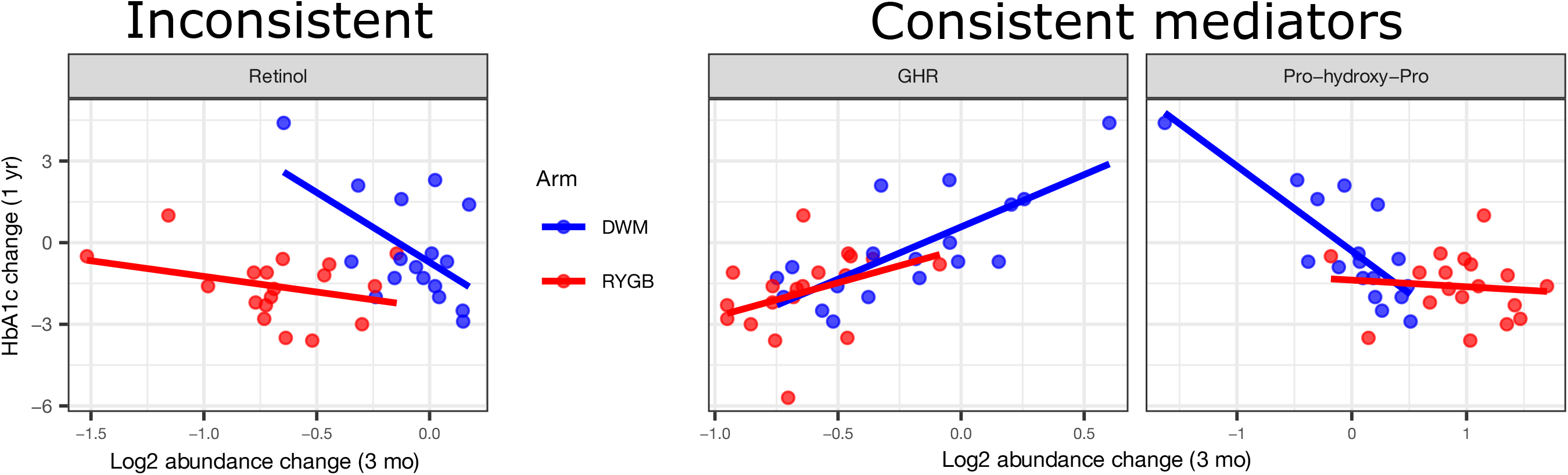
Mediation of HbA1c change at 1 year, see also Figure S1, S5, S6, Table S1, Table S2, and Text S1. X-axis represents log_2_ abundance change from baseline at the 3 month time point, and Y-axis represents HbA1c change from baseline at 1 year. Retinol is an inconsistent mediator, whereas GHR (growth hormone receptor) and pro-hydroxy-pro (prolylhydroxyproline) are consistent mediators of surgery’s effect.

A second limitation of the joint significance and other mediation methods is that they are intended for testing only one or a few mediators. Thus they do not include powerful statistical methods for high-throughput data, such as linear regression modeling with empirical Bayesian modeling of an analyte’s variance, which has been validated in multiple omics platforms, and is particularly powerful for small sample sizes (*44*).

To address these limitations, we developed a novel mediation method for high-throughput data, termed <u>Hi</u>gh-<u>t</u>hroughput <u>M</u>ediation <u>A</u>nalysis (Hitman). We demonstrate mathematically that it controls its false positive rate (**Text S1**), and it accounts for the direction of effect, so it does not identify inconsistent mediators as significant. For example, Hitman does not identify retinol as a significant mediator (p>0.9). Hitman applies empirical Bayesian linear modelling, so it has improved power in high-throughput settings (*44*).

Using simulations that followed the work of Barfield et al (*41*), we compared Hitman against the joint significance test and against a popular mediation test that uses the potential outcome framework (*45*) but was not included in previous simulations. We found that Hitman controls its false positive rate, like the other methods, and is significantly more powerful. For example, when the true causal mediation effects are of a size termed “small” (*40*), the joint significance and the potential outcome mediation test identify significance in <2.5% of simulations, whereas Hitman identifies >6%, which is significantly greater (p<10^-15^). When the true effect is of a size termed “medium” (*40*), the proportion of simulations identified as significant by the joint significance test and the potential outcome mediation test were 53.9% and 57.4%, respectively, whereas Hitman identified 68%, which is also significantly greater (p<10^-15^). These simulation results, simulation details, and a description of the Hitman algorithm are provided in **Text S1**.

Mediation methods can be misled by confounding variables, such as those that affect both the mediator and the outcome (*46*). To avoid confounding among measured mediators, some high-throughput mediation approaches decompose the measured analytes to identify latent, independent mediators (*47, 48*). However in biology many confounders may be unmeasured. For example surgery has multifaceted effects, so decomposing our measured analytes would not be sufficient to prevent the impact of confounders, so Hitman does not attempt to control for confounders. In biological datasets where all confounders and causal relationships between analytes cannot be controlled for, causal mediation analysis should be considered as exploratory (*49*).

### Proteomic, metabolomic, integrative pathway, and clinical mediation analysis

We applied Hitman to identify analytes whose early change (baseline to the 3 month time point) mediates HbA1c improvement at one year (**Table S2**). The only significant analyte was growth hormone receptor (GHR; p=10^-4^, FDR=0.12). The second-ranking analyte was CNDP1 (p=10^-3^). Both decreased >30% at the 3 month time point after RYGB (**Table S2**) and are shown in **Figure 6**. Robust GHR mediation led us to hypothesize that reductions in plasma GHR reflected reduced tissue content or altered receptor shedding, and thus could be associated with tissue-level growth hormone resistance. In the liver, GHR signaling typically increases secretion of the growth factor IGF1 (*50, 51*); however, IGF1 was not identified as a mediator by Hitman. Moreover, neither SOMAscan nor ELISA measures of IGF1 differed between arms at the 3 month time point (**Figure S5**), despite robust correlation between SOMAscan and ELISA changes (r=0.35, p=0.003). Even with unchanged IGF1 levels, plasma growth hormone levels were >6-fold higher in baseline-corrected RYGB vs. DWM at 12 months (**Figure S5**), raising the possibility that tissue- or pathway-selective growth hormone resistance in post-RYGB participants could contribute to sustained improvements in glucose metabolism (*52*).

The top-ranking putative metabolite mediators were prolylhydroxyproline (p<10^-3^, FDR=0.38) and isovalerylcarnitine (p<0.005; **Figure 4, Figure 6, Figure S4, Table S2**), which increased by 100% and decreased by 47% at the 3 month time point, respectively.

As a comparison to Hitman, we applied the joint significance method to test mediation of proteins and metabolites whose change at the 3 month time point mediates HbA1c improvement at one year. We found similar top analytes, but with weaker significance, consistent with our simulations demonstrating that Hitman offers substantial power (**Text S1**). The top-ranking analytes identified by the joint significance method were GHR (FDR=0.26) and pro-hydroxy-pro (FDR=0.73).

We next asked whether early postoperative change (baseline to the 3 month time point) in 40 clinical markers mediated HbA1c improvement at one year. The top-ranking early postoperative mediators identified by Hitman were 6 min walk test distance and BMI (p<0.03, FDR=0.4; **Table S2**). Strikingly, analytes identified in the 3 month time point mediation analysis (GHR and prolylhydroxyproline) were more significant (by p-value and FDR) than any clinical markers at the 3 month time point, suggesting the top analytes’ utility as potential clinical biomarkers.

We next sought to test mediation of our integrated pathways. There are several approaches that test pathway mediation (*48, 49*), but they are applied to the data itself, so they cannot take Hitman scores as input. However, the CAMERA pre-ranked procedure accepts our Hitman scores as input and accounts for correlation between genes (*53*). CAMERA tests pathway enrichment while accounting for correlation between genes. When we tested for metabolic pathways whose change at the 3 month time point mediate HbA1c improvement at 1 year, no pathways were significant. However, the top-ranking pathways were Beta-Alanine Metabolism (**Figure 5**) and Histidine Metabolism (p=0.006; FDR=0.23).

Given that improvements in glycemic control after RYGB are related to changes in insulin sensitivity and/or insulin secretion, we applied Hitman to identify analytes whose change from baseline at the 3 month time point mediated insulin secretion and insulin sensitivity change from baseline at 1 year (**Table S2**), and followed this with integrative pathway mediation analysis.

Insulin secretion, defined as the change in insulin from 0 to 30 minutes during a mixed meal tolerance test, was improved in RYGB vs. DWM (means of changes: RYGB=37.4, DWM=0.808; p=0.001). No single analytes were identified as mediators, but one pathway, Caffeine Metabolism, significantly mediated improved insulin secretion (FDR<10^-4^). Similarly, insulin sensitivity, defined by reduction in HOMA-IR, also improved in RYGB vs. DWM at 1 year (means of changes: RYGB=-2.12, DWM=-0.102; p=0.02). No individual proteins or metabolites were significant mediators of insulin sensitivity, but many top ranking analytes were BCAA-related metabolites. Consequently, pathway analysis identified Valine, Leucine and Isoleucine Degradation as a significant pathway mediator of insulin sensitivity (FDR<10^-7^), together with 14 other pathways, primarily involved in lipid and amino acid metabolism (**Table S2**).

## Discussion

We report findings from a clinical trial that randomized individuals with T2D to RYGB or medical management and measured high-throughput data serially over 3 years. This design allowed us to identify analytes and pathways with differential patterns of change and potential mediators of improved metabolism after RYGB, providing candidates for nonsurgical approaches to T2D. This is shown in a graphical overview in **Figure S6**. As expected, RYGB exerted greater weight loss and glycemic improvement, and observed protein and metabolite changes are more robust during periods of active weight loss, indicating partial weight dependence of metabolic effects. However, differential abundance of many analytes remained after BMI adjustment, indicating weight-independence. Moreover, the top analytes at the 3 month time point are more significant mediators of glycemia at 1 year than any clinical markers, including BMI, indicating that these analytes may have clinical significance as biomarkers of bariatric surgery or therapeutic targets. Our mediation analysis tool, Hitman, provides a new method to analyze studies that randomize a treatment and observe differences in outcomes; using measurements of analytes upstream of the outcome, putative mediators of the outcome can be identified.

One feature of Hitman that improves its power is that it accounts for the direction of mediation. Hitman finds that retinol’s direction of mediation is inconsistent with mediation of improved glycemic control after RYGB. As seen in **Figure 3**, RYGB reduces retinol more than DWM, which is compatible with previous reports in bariatric surgery (*54–56*). However, within both treatment groups, greater retinol reduction was associated with increased HbA1c, indicating that the reduction in retinol is not a likely mediator of improved glycemia. Similarly, retinol binding protein 4 (RBP4) was numerically lower after RYGB, consistent with prior reports linking increased serum RBP4 levels with obesity and insulin resistance (*57*). Like retinol, RBP4 reduction was associated with increased HbA1c within each of the treatment groups, so it was also penalized by Hitman (**Table S2**). Thus, these data highlight complex relationships between retinol, RBP4, and T2D phenotypes, but suggest that these changes are not likely to mediate improved glycemia after RYGB.

Several significant pathways identified were driven by reductions in CNDP1, a top-ranking putative protein mediator of HbA1c. CNDP1 is a secreted dipeptidase that hydrolyzes carnosine to β-alanine and L-histidine. CNDP1 was found to decline significantly 3 months after bariatric surgery (*58*). These relationships are consistent with a common *CNDP1* genetic variant that enhances enzymatic activity, which is associated with loss of glycemic control in mice (*59*).

The significant protein putative mediator of improved glycemic control was GHR, which was decreased in plasma from post-RYGB participants. Interestingly, reduction in GHR expression was recently demonstrated in multiple tissues of post-RYGB rodents (*60*). Parallel reductions in GHR and increased growth hormone levels post-RYGB suggest the possibility that tissue-selective growth hormone resistance could contribute to improvements in insulin sensitivity and reductions in hepatic glucose production, as observed in humans treated with a growth hormone antagonist (*52*).

The top putative metabolite mediator of HbA1c was prolylhydroxyproline. Prolylhydroxyproline is a marker of collagen degradation, and the increased levels in RYGB could be linked to increased bone turnover (*61*) consistently observed after bariatric surgery, including in the present cohort (*62, 63*). One possible mechanism for mediation is that prolylhydroxyproline can facilitate adipose-derived stromal vascular cells to differentiate into more metabolically active beige adipocytes (*64*).

Valine, Leucine, and Isoleucine Degradation was a significant HOMA-IR pathway mediator and the most significantly changed pathway at 3 months between groups. There were robust post-RYGB decreases in BCAA and multiple downstream catabolic intermediates, including C3 and C5 acylcarnitines, with isovalerylcarnitine as a putative causal metabolite mediator. Our results are consistent with prior findings of increased BCAA in insulin resistance (*65, 66*) and reduced BCAA in response to RYGB in nonrandomized studies (*25–27*). Lower BCAA levels post-RYGB may be a consequence of weight loss-linked improvements in insulin sensitivity or altered microbial metabolism (*34*), but may also contribute directly to improved insulin sensitivity (*67*). Our data reveal a marked 88% reduction in the valine catabolic intermediate 3-hydroxyisobutyrate at the 3 month time point post-RYGB. This is particularly interesting since 3-hydroxyisobutyrate can exit mitochondria and serve as a signaling molecule, promoting muscle lipid uptake and insulin resistance (*68*) and impaired mitochondrial OXPHOS activity (*69*).

We acknowledge that profiling semi-quantitative plasma metabolomics and proteomics, with emphasis on the secreted proteome, cannot fully define the pleiotropic effects of bariatric surgery, including both weight-dependent and weight-independent changes in complex interorgan communication, gut microbiome effects, or tissue-specific flux in metabolic pathways. Our study is innovative as we have developed and applied novel bioinformatics tools to identify clinical measures, analytes and pathways that mediate response to RYGB within a randomized clinical trial comparing RYGB vs. medical management in T2D. The analytes we have identified and validated can be modulated in future studies to determine whether they can be utilized for non-surgical control of glucose metabolism in T2D.

## Materials and Methods

### Clinical study

Participants were randomized to RYGB, performed using standard operative protocols, or DWM, conducted by a multidisciplinary team in groups of 10-15 via the Why WAIT (Weight Achievement and Intensive Treatment) program (*70*). Both RYGB and DWM groups returned to usual care following intervention, and annual follow-ups after one year were observational. The protocol was approved by the Partners Healthcare Institutional Review Board, and an independent data monitoring committee reviewed patient safety. Difference of outcomes between groups was analyzed with a t-test, and significance was defined as p<0.05.

### Metabolic assessments

Metabolic assessments were performed at baseline and repeated at the 3 month time point, which was defined by achieving 10% of initial body weight loss or 3 months, if 10% weight loss had not yet been achieved by this time point. This time point was defined to permit assessments at a similar level of weight loss in both groups. Assessments were also repeated in both groups at 12, 18, 24, and 36 months. Analysis of blood samples collected in the fasting state included HbA1c, plasma glucose, and lipids (Quest Diagnostics). Aliquots were stored at −80 C.

### Proteomic profiling and validation

Plasma proteome profiling was performed using the high-throughput DNA aptamer-based SOMAscan assay platform (SomaLogic, Inc.)(*71*). Abundance of 1129 proteins (enriched for extracellular proteins) was quantified as relative fluorescent units (RFU), normalized, calibrated, and log_2_-transformed. Samples were available from 38 participants at baseline and 38, 35, 25 and 23 participants at the 3, 12, 24, and 36 month time points, respectively (**Table S1**).

Selected proteomic data were validated in a subset of fasting plasma samples using specific ELISA, including IGFBP2 (22-BP2HU-E01, ALPCO, NH), CNDP1 (F34010, LifeSpan Biosciences, WA), growth hormone (DGH00, R&D Systems, MN), and total IGF-1 (DG100, R&D Systems, MN). For RBP4 and TTR, plasma samples were assayed using quantitative western blotting using a polyclonal antibody to human RBP4 (Dako) and human TTR (Dako) with standard curves of purified human RBP4 (*72*) or TTR (Sigma) protein on each blot (*57*). Changes per individual over time were tested for positive correlation to corresponding SOMAscan changes with a one-sided test of Pearson correlation.

### Metabolomic profiling

Plasma metabolomics were profiled using a commercial semi-quantitative mass spectrometrybased platform (Metabolon, Inc.)(*73*). Metabolites that had missing values in more than 85% of samples were filtered out, missing values were imputed with half of the minimum for each metabolite, and abundance values were log_2_-transformed. Samples were available from 38 participants at baseline and 36, 35 and 22 participants at 3, 12, and 36 months; due to cost, metabolomics were not profiled at 24 months.

### Differential abundance of proteomics, metabolomics, and pathways

To test differential abundance of log_2_ normalized analytes between groups at baseline, we applied moderated t-tests with our R package ezlimma, which streamlines and extends the R package limma (*44, 74*). Limma applies linear regression modeling with empirical Bayesian methods to improve each analyte’s variance estimation using analytes’ shared systematic variance. At post-baseline time points, we calculated change in analyte abundance from baseline for each individual, and then applied moderated t-tests to test if these changes varied by group. We repeated these post-baseline analyses accounting for each person’s BMI change. We similarly applied limma’s roast method (*75*) for differentially abundant pathways via our ezlimma package. We considered Roast’s “Mixed” statistics, which correspond to assessing the absolute value of an analyte’s change, so that if some of a pathway’s analytes are upregulated (e.g. proteins) and others are down-regulated (e.g. metabolites), these changes do not cancel out the pathway’s effect. Significance for this and other omics analyses was defined as FDR<0.15. In plots of individual analytes with standard error of the mean (SEM), ordinary (i.e. unmoderated) SEM are displayed.

### Mediation analysis

We tested mediation of each clinical variable with the causal chain: group → clinical variable change → clinical outcome change. We tested each clinical variable’s mediation by defining *group* as a binary variable representing RYGB or DWM per individual; *clinical variable change* as each clinical variable’s change between baseline and the 3 month time point per individual; and *clinical outcome change* as the change in clinical outcome between baseline and 12 months per individual.

We tested mediation of analytes with the causal chain: group → analyte change → clinical outcome change. We tested each analyte’s mediation by defining *group* as a binary variable representing RYGB or DWM per individual; *analyte change* as each analyte’s change (on the log_2_ scale) between baseline and the 3 month time point per individual; and *clinical outcome change* as the change in clinical outcome between baseline and 12 months per individual. As a comparison to Hitman, we similarly tested mediation of analytes using the joint significance method and using the mediation package (*45*) in the R software. Simulation results were compared statistically with a two-sample t-test with pooled variance in the R software.

We tested these Hitman results against pathways containing proteins and metabolites from the Small Molecule Pathway Database (SMPDB)(*37*) with the CAMERA pre-ranked pathway analysis method (*53*) from the Limma package.

### Data and software availability

SOMAscan and clinical data have been deposited at GEO:GSE122279. SOMAscan, metabolomics, and clinical data, and the R/Bioconductor (*74*) code to reproduce main results from them (including Table S2 and pathway results with links to the underlying pathway’s analyte statistics) are available at https://github.com/jdreyf/slimm-t2d-omics. Our streamlined limma R package ezlimma is available at https://github.com/jdreyf/ezlimma. The Hitman package is available at https://github.com/jdreyf/Hitman.

## Supplemental Information Items

**Figure S1.**
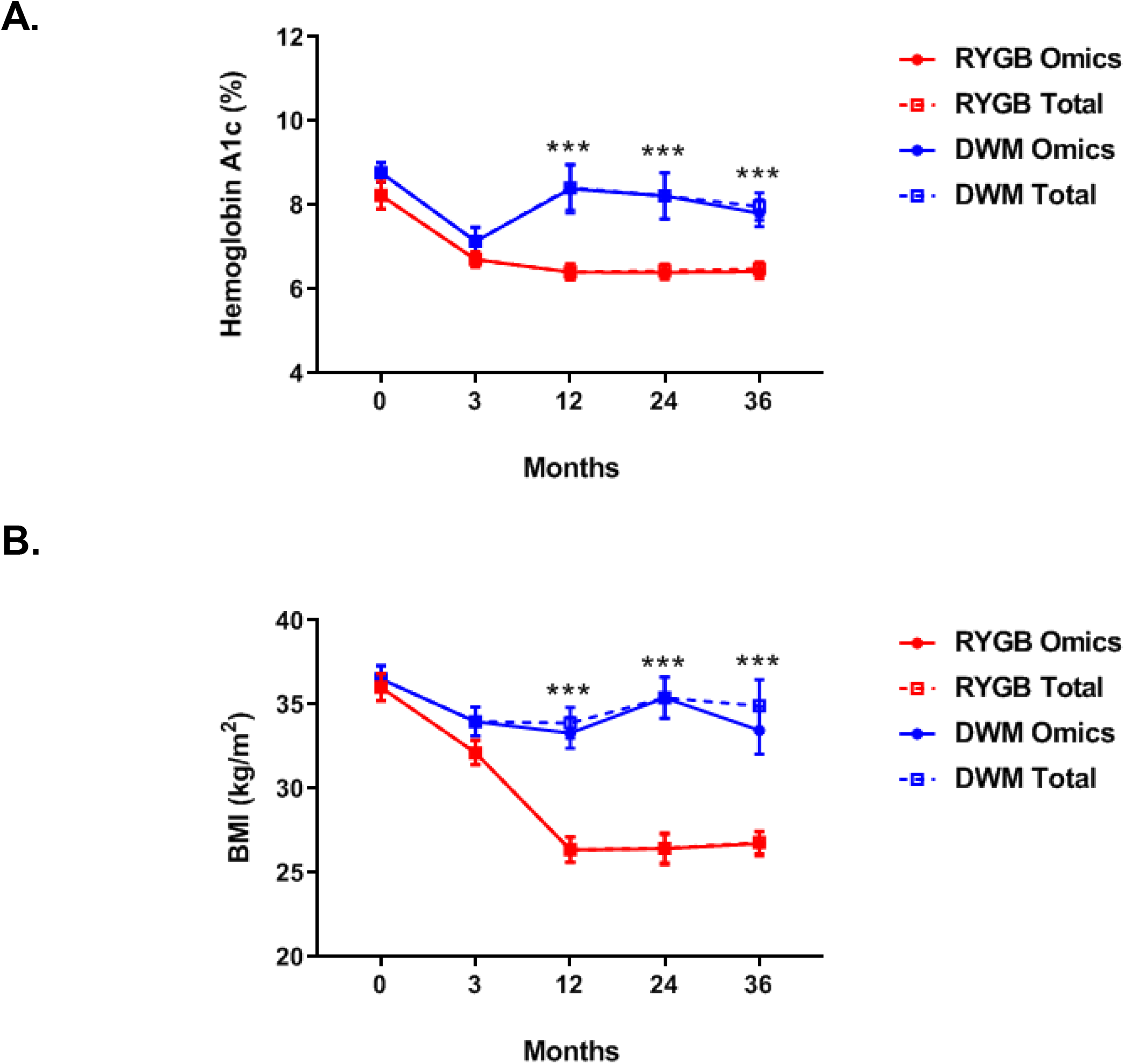
HbA1c and BMI for SLIMM-T2D cohort and those that had omics per time point. (A) HbA1c and (B) BMI in both treatment arms for the SLIMM-T2D cohort (“Total”) and those that had proteomics or metabolomics per time point (“Omics”). Data are reported as mean ± SEM. Data were analyzed by t-tests. For HbA1c and BMI, no significant difference was found between the Total and Omics groups within RYGB or within DWM at any time point. However, differences between RYGB and DWM per time point were found: *p<0.05, **p<0.01, ***p<0.001 and #p<0.0001.

**Figure S2.**
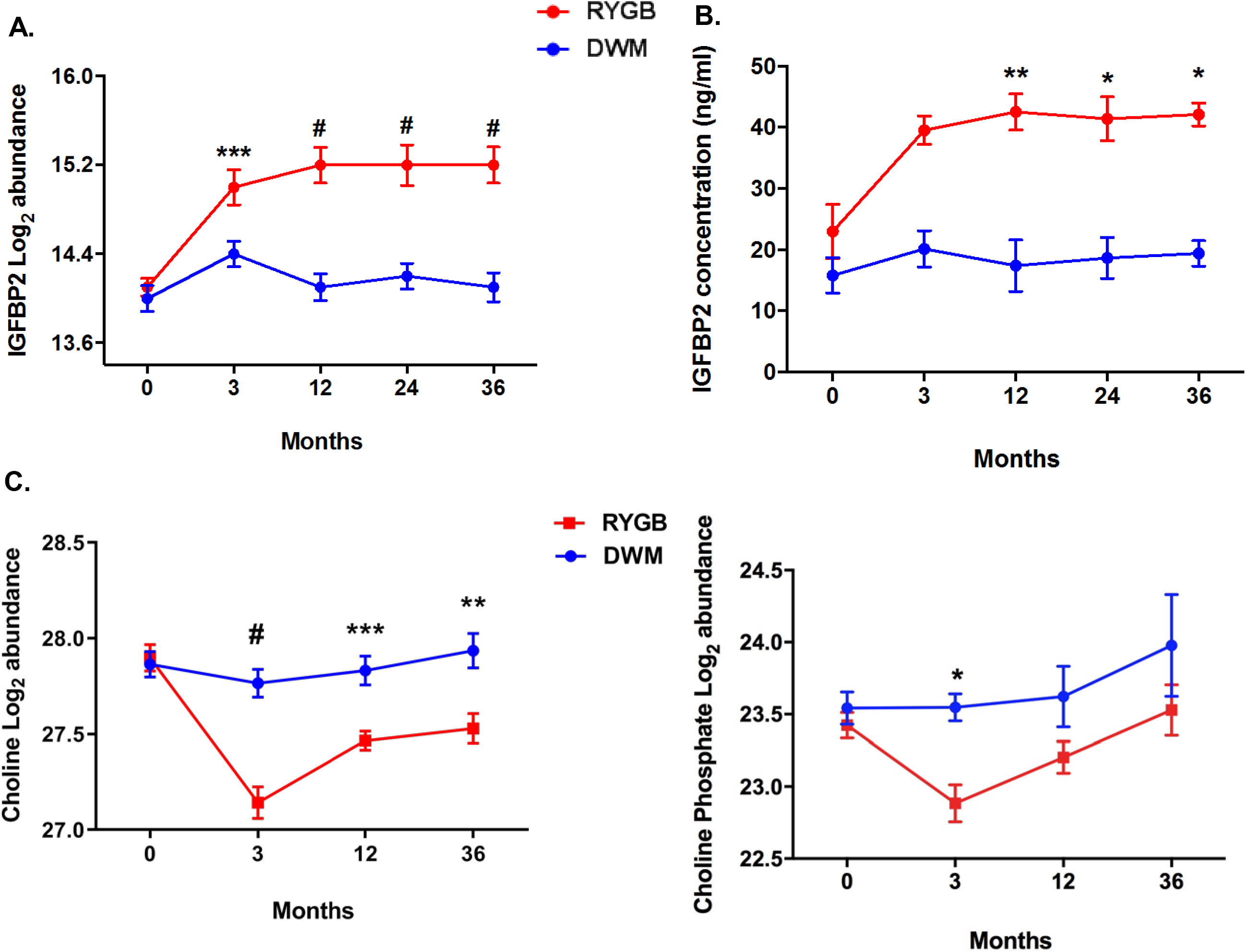
IGFBP2, choline, and choline phosphate levels. (A) IGFBP2 log_2_ abundance measured by SOMAscan. (B) IGFBP2 concentration measured by ELISA. (C-D) Choline and choline phosphate log_2_ abundance measured by metabolomics. Data were analyzed by moderated t-tests; post-baseline time points were baseline-corrected. Data are reported as mean ± SEM: *p<0.05, **p<0.01, ***p<0.001, and #p<0.0001.

**Figure S3.**
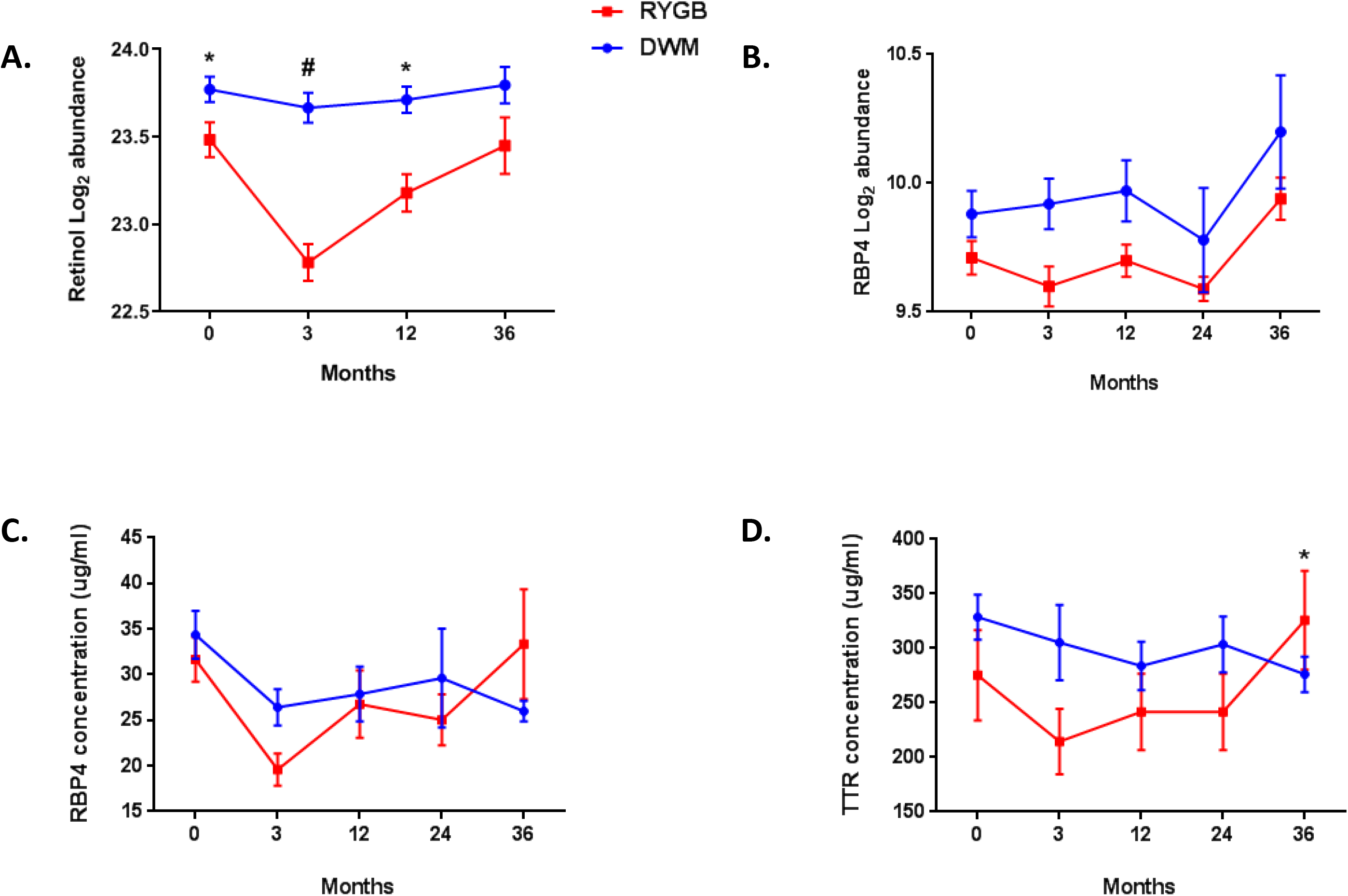
Retinol, RBP4, and TTR abundance. (A) Log_2_ abundance of retinol measured by metabolomics. (B) Log_2_ abundance of RBP4 measured by SOMAscan. (C) RBP4 and (D) TTR plasma levels measured by quantitative western blot. Data were analyzed by moderated t-tests; post-baseline time points were baseline-corrected. Data are reported as mean ± SEM: *p<0.05, **p<0.01, ***p<0.001 and #p<0.0001.

**Figure S4.**
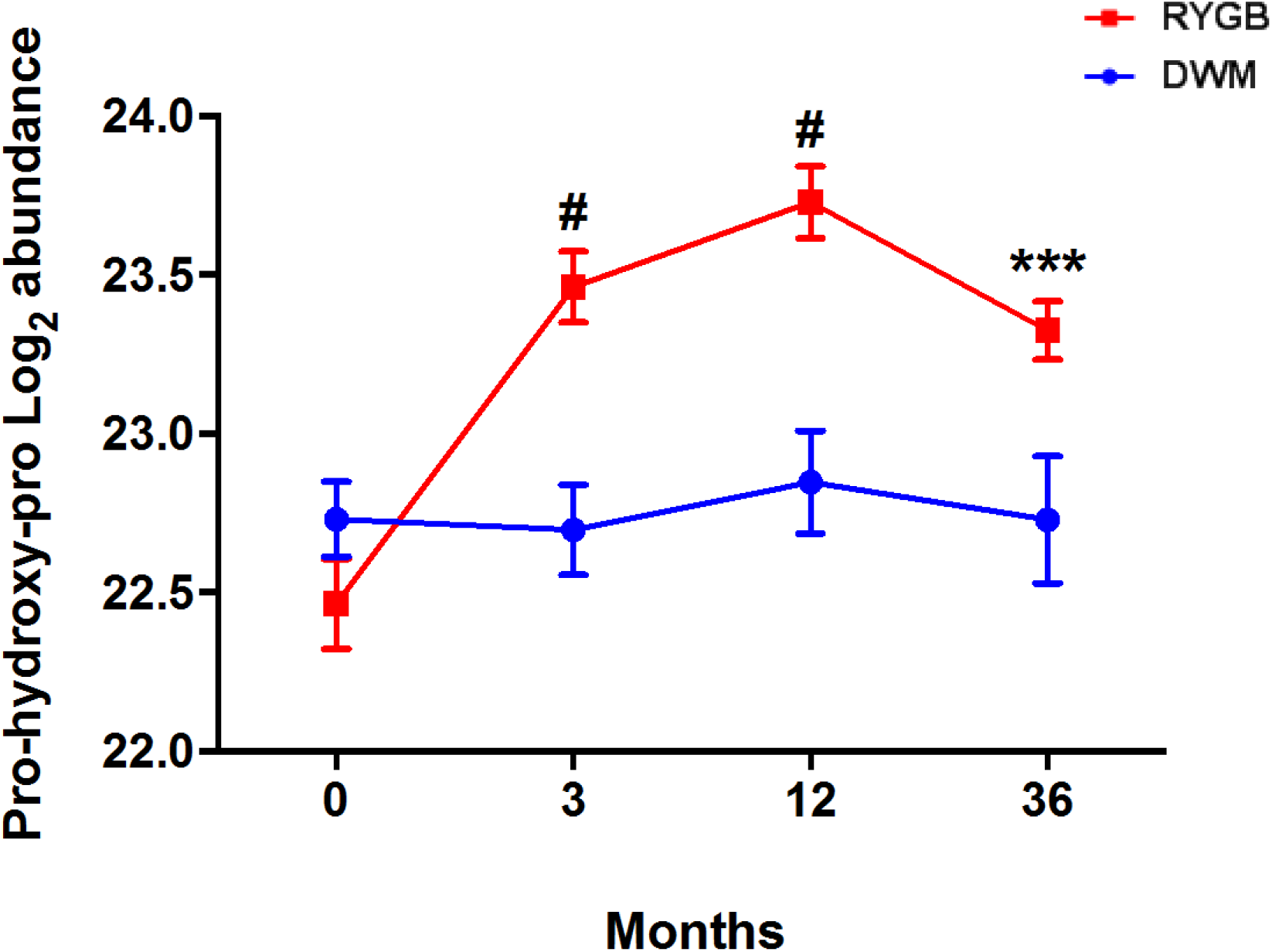
Abundance of prolylhydroxyproline over time. Log_2_ abundance of the mediator prolylhydroxyproline measured by metabolomics. Data were analyzed by moderated t-tests; post-baseline time points were baseline-corrected. Data are reported as mean ± SEM: *p<0.05, **p<0.01, ***p<0.001 and #p<0.0001.

**Figure S5.**
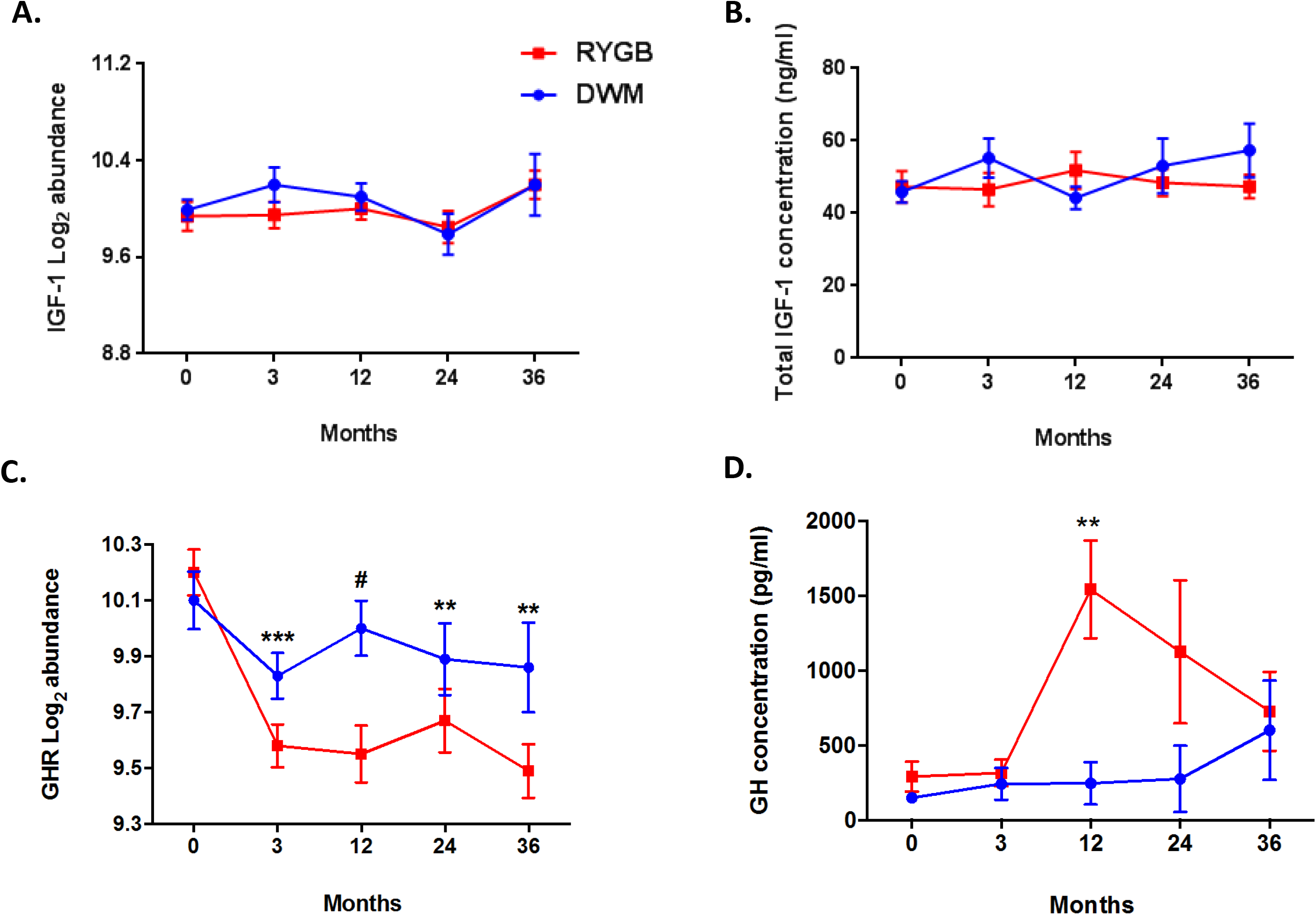
IGF-1 and GHR SOMAscan and ELISA levels. (A) Log_2_ abundance of IGF-1, measured by SOMAscan. (B) Plasma levels of total IGF-1, measured by ELISA. (C) Log_2_ abundance of GHR, measured by SOMAscan. (D) Plasma levels of growth hormone, measured by ELISA. Data were analyzed by moderated t-tests; post-baseline time points were baseline-corrected. Data are reported as mean ± SEM: *p<0.05, **p<0.01, ***p<0.001 and #p<0.0001.

**Figure S6.**
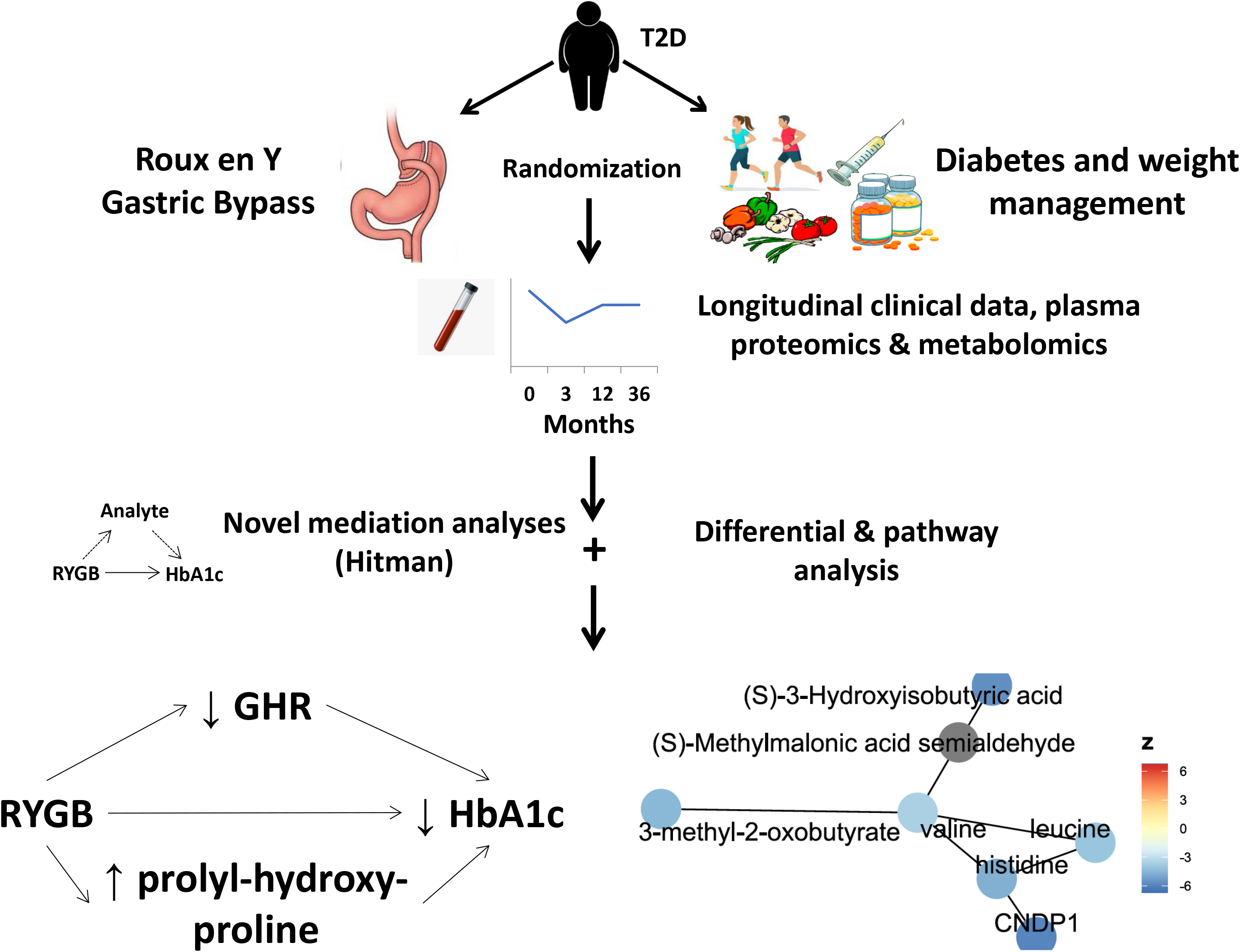
Graphical overview. Shows study flow. Bottom show top two mediators (GHR and prolyl-hydroxy-proline) and a network composed of top analytes from Valine, Leucine, and Isoleucine Degradation and Beta-Alanine Metabolism. Nodes colored dark grey were not measured here.

**Table S1. Baseline demographics of participants that had both proteomics and metabolomics at all of baseline, 3 months, and 12 months; and number of participants taking each diabetes medication class per arm at each time point.** The metabolic characteristics of the subset of participants who had omics data did not differ between groups at baseline. There were differences in medication use between groups.

**Table S2. Differential abundance, correlation, and mediation tables.** Differential abundance of proteins, metabolites, and pathways; Pearson correlation coefficients of top proteins vs. top metabolites; mediation of HbA1c, insulin secretion (as the change in insulin from 0 to 30 minutes during a mixed meal tolerance test), and HOMA-IR.

**Text S1. Hitman algorithm and validation.** Technical description and validation.

## Acknowledgements

We thank Danyel Cavazos for help with figures; Grace Daher, G. H. Nguyen, Ronaldo Da Silva Francisco Junior, and Lado Gholijashvili for assistance with R package development; Vera Djordjilovic for helpful discussion on Hitman; and Harvard Medical School Research Computing for computational resources.

## Trial Registration

Clinicaltrials.gov NCT01073020

## Text S1. High-throughput mediation analysis (Hitman)

### Model

Our causal model of the effect of the treatment or exposure (*E*) on the outcome (*Y*), and the effect’s mediators (*M*) are shown in Figure 1a and 1b. We have *E* → **M** → *Y* and *E* → *Y*, where arrows (→) represent potential directed causal effects, with structural parameters next to the arrows. Our unmediated model is 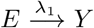, where the structural parameter *λ*_1_ represents the *total effect* (http://davidakenny.net/cm/mediate.htm). Our causal mediation model per mediator is shown in Figure 1b. We assume that *E* causally affects *Y*, and that this total effect is robust. We are then interested in the mediation effect (or *indirect effect*), represented by *α*_1_ *θ*_2_. *θ*_1_ is the *direct effect* per mediator. Lack of an arrow indicates no causal effect (Pearl 2010), so the model assumes that neither *M* nor *Y* affect *E* (as occurs when *E* is randomized), and that *Y* does not affect *M*. Hitman does not test these assumptions.

**Figure 1:**
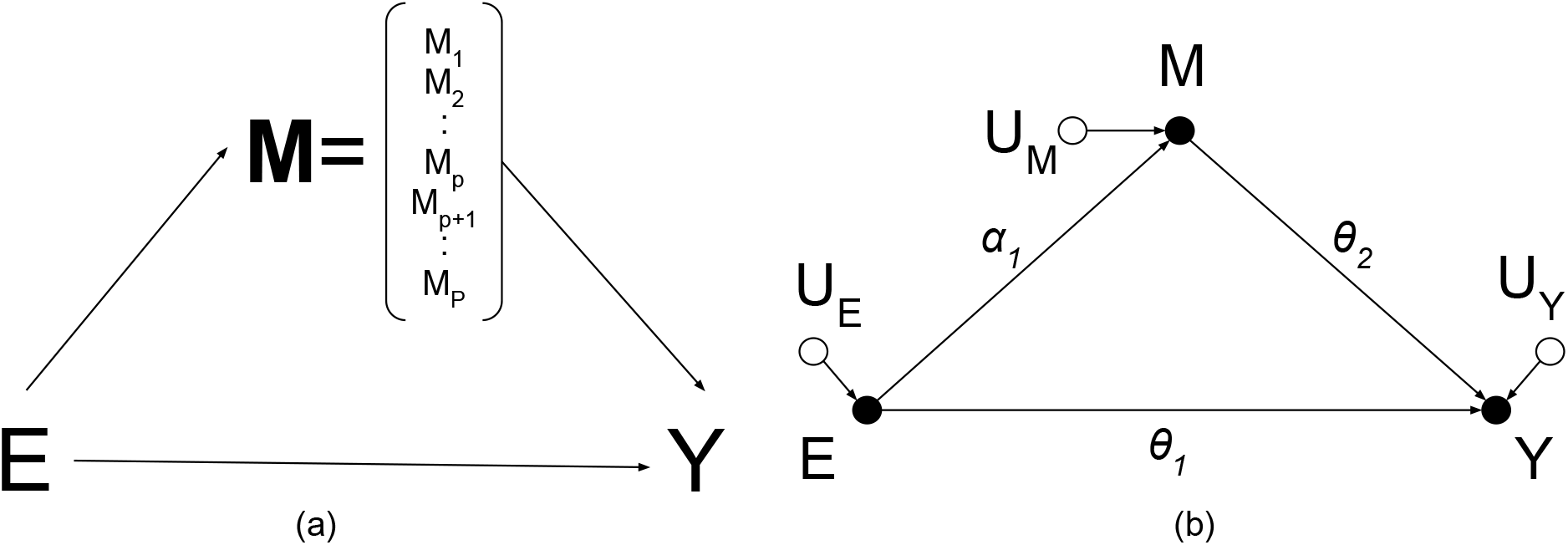
Hitman model without covariates.

Our model in Figure 1a is based on a biological system, such as a cell, where an intervention (or exposure) like a gene knockout can have effects across all activities in the cell. The number of potential mediators of the exposure is *P*, where *P* is astronomically large. In an omics experiment, we measure *p* (with *p* much smaller than *P, p* ≪ *P*) analytes across *n* samples, with *p* >> *n*. The *P* mediators in Figure 1a have unknown relationships among themselves and on the outcome. Ideally, we would identify these relationships so that we can account for them, but with few samples and an astronomical number of mostly unmeasured mediators, we do not attempt that here, and instead pursue an exploratory, one-at-a-time analysis of our measured mediators, so that we can identify which mediators to pursue with further studies.

Our linear structural equation model (SEM) per mediator with covariate vector or matrix *X* is:

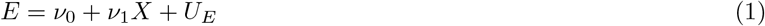

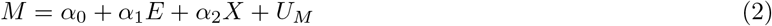

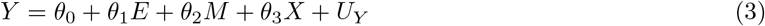

where *U* terms represent omitted factors that explain sources of variation. The SEM’s coefficients represent structural parameters estimable from the data. These terms must be mutually independent for the validity of mediation analyses. When we randomize *E*, we know that *UE* is independent of the other *U*’s. However, *U_M_* is often dependent with *U_E_*, due to another mediator (likely unmeasured) affecting both *M* and *Y*. So, “theoretical knowledge must be invoked to identify the sources of these correlations and control for common causes (so called ‘confounders’) of *M* and *Y* whenever they are observable” (Pearl 2014a). Such confounders can be included in *X*. When *X* is a matrix, *ν*_1_, *α*_2_, and *θ*_3_ are vectors.

We implement Hitman for our data with the assumptions that *E* is randomized and, under the null, the *U*’s are independent given *X* and *U_M_* and *U_Y_* have known distributions. For example, our analyte mediators and outcome follow the normal distribution. Hitman also accommodates outcomes that follow any generalized linear model, such as logistic regression for binary *Y*.

Hitman is based on the joint significance or causal steps test. The joint significance test requires that the exposure causes the outcome (Judd & Kenny 1981)(MacKinnon et al. 2002), and has been shown to have more power than the product method and to control its false positive rate, because it is an intersection-union test (Huang et al. 2018). A drawback of the joint significance test is that it doesn’t account for direction of effect. For example, if *E* increases *Y* (i.e. an increase in *E* increases *Y*), then we would want mediators *M* where *E* increases *M* and *M* increases *Y*, or where *E* decreases *M* and *M* decreases *Y*. However, the joint significance test might find as significant an “inconsistent” mediator where *E* increases *M* and *M* decreases *Y* (MacKinnon et al. 2002). We address this in Hitman.

### Method

For each mediator (*M*), the null hypothesis is that *E* has no effect on *M* (*α*_1_ = 0); or that *M* has no effect on *Y* (*θ*_2_ = 0); or that the direction of mediation, sgn(*α*_1_ *θ*_2_), is not consistent with the direction of the total effect, sgn(*λ*_1_), with *sgn* being the sign function. An example of an inconsistent mediator is *E* increases *Y*, but *E* increases *M* and *M* decreases *Y*. The alternative hypothesis is that *E* affects *M* and *M* affects *Y*, and that the direction of *E* → *M* and *M* → *Y* are consistent with that of the total effect of *E* on *Y*. This tests if *M* explains at least some of the dependence between *E* and *Y*; *M* need not explain all of it.

Hitman implements tests with high-throughput mediators using the R/Bioconductor linear modeling package limma (Ritchie et al. 2015). Limma models the variance of features (e.g. proteins or metabolites) with an empirical Bayesian method, which exploits information about shared technical variance between features for improved power, especially when the sample size is small (Ritchie et al. 2015). Limma can also account for a mean-variance relationship, as occurs in RNA-seq (Law et al. 2014). To model variance of feature abundance in linear modeling, limma models feature abundance as the dependent variable.

To estimate the direction of mediation, Hitman uses sgn 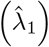 from the linear model with covariates *Y* = *λ*_0_ + *λ*_1_*E* + *λ*_2_*X* + *ϵ*. This model does not include mediators, because here we are not interested in the indirect or mediated effects. Hitman assumes the total effect is robust, so treats the estimated direction as correct, i.e. 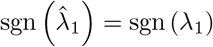.

For each mediator *M*:

1. Measure effect of *E* and *M* using equation 2. Here and later, if *X* is absent, remove its term. If *X* has multiple columns, *α*_2_ is a vector. Test *α*_1_ = 0 in limma and define the resulting p-value *p*_1_.
2. Measure association of *M* and *Y* given *E* (and possibly *X*) using equation 3. We would like to test *θ*_2_ = 0 using limma. To model the variance of *M* with empirical Bayesian methods, we need to make *M* the dependent variable. So we use an approach similar to partial correlation.

i. Estimate the residuals of *Y* = *λ*_0_ + *λ*_1_*E* + *λ*_2_*X* + *e* as *e_Y_*.
ii. Estimate the residuals of *M* = *α*_0_ + *α*_1_*E* + *α*_2_*X* + *e* as *e_M_*.
iii. From the linear model *e_M_*=*θ*_2_*e_Y_* + *ϵ*, test *θ*_2_ = 0 in limma and define the resulting p-value *p*_2_. Now define

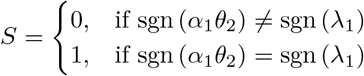

and

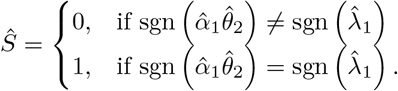
3. If *Ŝ* = 1, then the direction of total effect agrees with that of *E* → *M* → *Y*, and the final p-value = 0.5max(*p*_1_, *p*_2_). Otherwise, the direction of the indirect effect is inconsistent with that of the overall effect, and the final p-value = 1 – 0.5max(*p*_1_, *p*_2_). The intuition behind the test is that *Ŝ* allows for one-sided testing of 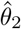 or 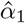.
4. To account for testing multiple mediators, calculate false discovery rates (FDRs) from the mediator p-values (Benjamini & Hochberg 1995).

### Example

We illustrate Hitman through an example of an inconsistent mediator (Figure 2). This figure is reminiscent of Simpson’s Paradox (Pearl 2014b). The exposure increases the outcome, and it increases the mediator. Moreover, the mediator is associated with the outcome, but it *decreases* the outcome, resulting in inconsistent mediation.

**Figure 2:**
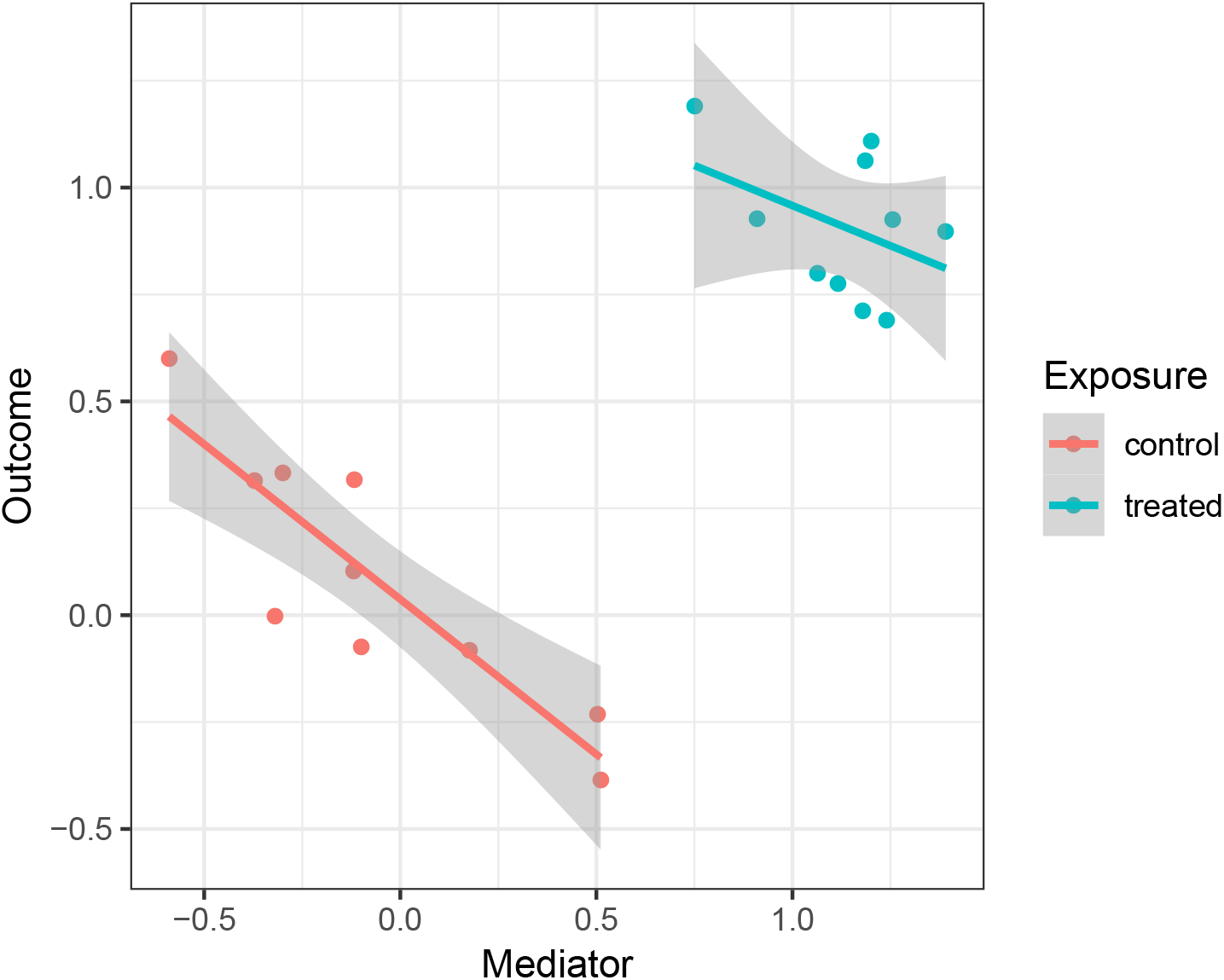
Inconsistent mediator.

To define this generic example, let the realized value of the random estimate 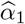 be *a*, and similarly the realized value of 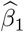 be *b*, and of 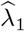 be *g*. In this example, *a*>0, *g*>0, and *b*<0. Then *sign*(*g*) ≠ *sign*(*ab*). However, this evidence is opposite the alternative hypothesis, where we want the direction of effect to be consistent: sgn(*λ*_1_) = *sign*(*α*_1_*β*_1_). Thus, Hitman conservatively estimates this mediator’s p-value as 1, whereas if consistency was not accounted for, the p-value would be < 0.001.

### Mathematical justification

We show here that Hitman properly controls its false positive rate in theory. Below we show this in simulations. Hitman, like the joint significance method, requires that the exposure causes the outcome. Hitman also requires that this effect is robust, so the sign of the estimated total effect matches the sign of the total effect.

Then for a single mediator, the null and alternative hypotheses as described above are:

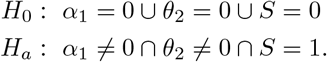

But the terms in *H_a_* are not independent, because, e.g. *S* = 1 → *α*_1_ ≠ 0 ⋂ *θ*_2_ ≠ 0, and consequently we could also write *H_a_*: *S* = 1.

To show that Hitman controls its size given the composite null, we consider the most challenging case under *H*_0_. Without loss of generality, under the assumption that the estimated direction is correct, let 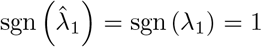. Hitman treats *α*_1_ and *θ*_2_ symmetrically, so without loss of generality we select *θ*_2_ to have a defined sign, and without loss of generality consider sgn(*θ*_2_) = 1. For the case to be maximally challenging: *θ*_2_ → ∞, so 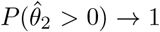 and *p*_2_ → 0. Under the null, we have *S* = 0, and *S* = 0 → *α*_1_ ≤ 0, so we consider *α*_1_ = 0, because this is the most challenging value for Hitman.

The p-value from Hitman can be defined as a mixture for this challenging case:

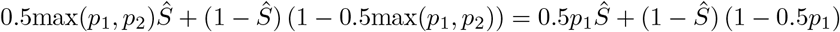

To demonstrate this is a valid p-value, we need to show *P*(*p* ≤ *u*|*H*_0_) ≤ *u* for each 0 ≤ *u* ≤ 1. We have that

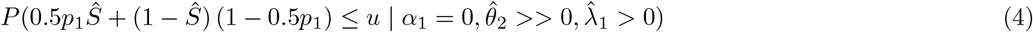

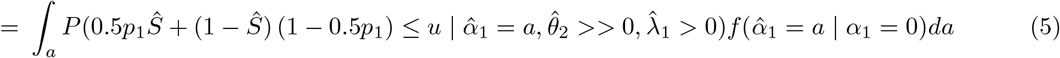

where ≫ denotes much larger than; we leave *S* = 0 out of equations 4 and 5, because it is implied by the other two parameters; and equation 5 follows from equation 4 by the law of total probability. We have that 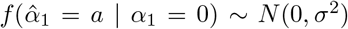. Without loss of generality, we conveniently assume *σ* = 1, so 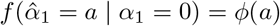 and 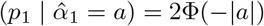, because 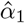 is tested with a two-sided test, where *ϕ* is the standard normal density and Φ is the standard normal cumulative distribution function. So, equation 5 simplifies to

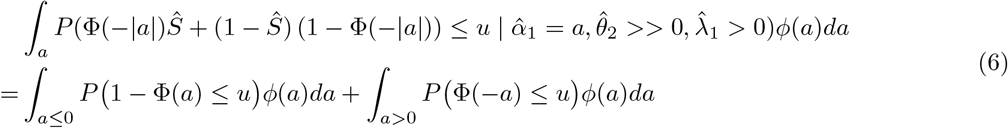

Because if *a* ≤ 0, then *Ŝ* = 0 and Φ(–|*a*|) = Φ(*a*), whereas if *a* > 0 then Φ(–|*a*|) = Φ(–*a*). We further simplify the second term in equation 6:

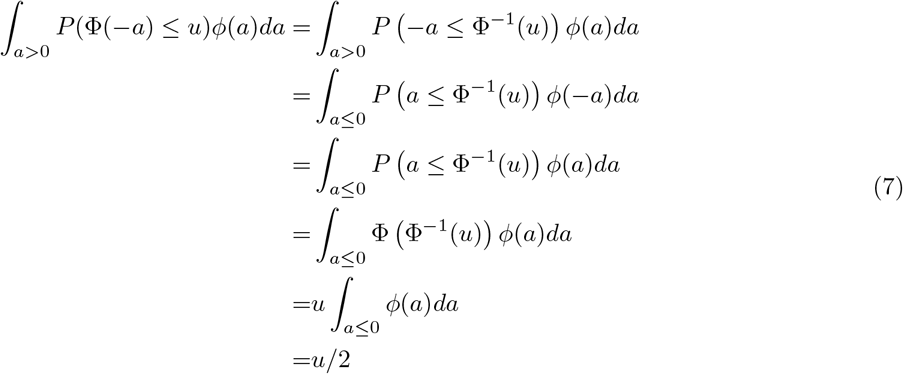

We also simplify the first term in equation 6:

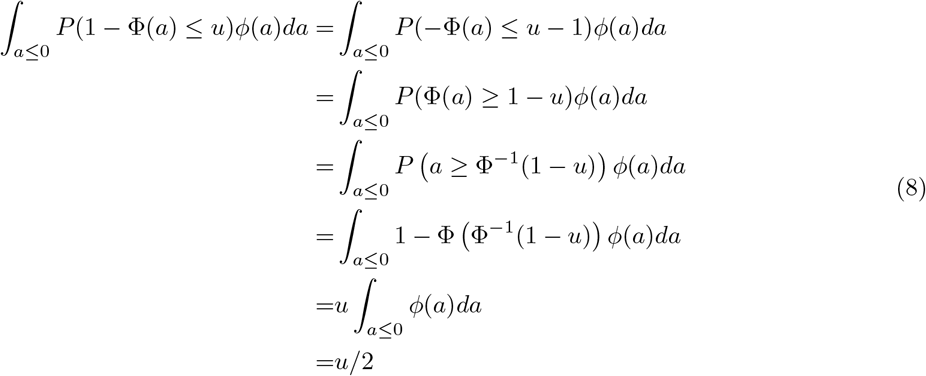

Thus, ∀*u*: 0 ≤ *u* ≤ 1, *P*(*p* ≤ *u* | *H*_0_) ≤ *u*, so Hitman controls its false positive rate.

### Simulations

We validated our size and power following a simulation study (Barfield et al. 2017). This study’s parameters were a subset of a previous simulation study (MacKinnon et al., 2002). Like (Barfield et al. 2017), we simulated data from *Y* = *t*_0_ + *t*_1_*E* + *t*_2_*M*_1_ + *t*_3_*X* + *e_Y_*, where *M*_1_ defines the first mediator, which was simulated as *M*_1_ = *b*_0_ + *b*_1_*E* + *b*_2_ *X* + *e*_*M*_1__. This was the only mediator associated with exposure and outcome, and so was the only mediator whose p-value we examine.

Hitman relies on high-throughput data, so we simulated other mediators as *M_i_* = *b*_0_ + *b*_2_*X* + *e_M_i__* for *i* = 1, which are independent of the exposure and the outcome. (We index mediators using the subscript *i*, but suppress indices that indicate sample.) These other mediators are exploited by limma to estimate the shared technical variance, as would happen in an omics study.

*X* and *E* and the error terms *e_Y_* and *e_M_i__* for all *i* were simulated as independent standard normal variables. We set *θ*_0_ = *θ*_3_ = *β*_0_ = *β*_2_ = 0.14. Barfield et al. (2017) set the direct effect to be “small”, *θ*_1_ = 0.14.

Hitman is only applicable when there is a robust overall effect, but these simulations include cases where there is no overall effect. Thus these simulations are not entirely appropriate for Hitman. However, we addressed this in part by setting the direct effect *θ*_1_ to be “large” (*θ*_1_ = 0.59) as per (MacKinnon et al., 2002). In this way, the simulations more closely match Hitman cases but can still be compared to previous simulations (Barfield et al., 2017; MacKinnon et al., 2002).

Like Barfield et al. (2017), we simulated all combinations of *θ*_2_, *β*_1_ ∈ (0, 0.14, 0.39), which correspond to effects of “zero”, “small,” and “medium” size, respectively (MacKinnon et al., 2002). We tested in what proportion of 10,000 simulations *M*_1_ of 100 mediators across 50 samples achieved a p-value ≤ 0.05. Our results for these parameter values for Hitman, the joint significance test, and the *mediate* function from the R package *mediation* are shown in Table 1.

**Table 1:** Comparison of methods via simulation.

| $\beta_1$ | $\theta_1$ | Hitman | joint | mediate |
| --- | --- | --- | --- | --- |
| 0.00 | 0.00 | 0.006 | 0.002 | 0.002 |
| 0.00 | 0.14 | 0.012 | 0.006 | 0.005 |
| 0.14 | 0.00 | 0.016 | 0.008 | 0.007 |
| 0.00 | 0.39 | 0.040 | 0.036 | 0.030 |
| 0.39 | 0.00 | 0.048 | 0.038 | 0.028 |
| 0.14 | 0.14 | 0.063 | 0.023 | 0.015 |
| 0.14 | 0.39 | 0.206 | 0.108 | 0.103 |
| 0.39 | 0.14 | 0.207 | 0.111 | 0.127 |
| 0.39 | 0.39 | 0.682 | 0.539 | 0.574 |

A statistical test’s size is the probability of falsely rejecting the null hypothesis, which is the probability of a false positive or a Type 1 error. All the methods here control their size, as they maintain a false positive rate less than 5%.

A statistical test’s power is the probability that the test correctly rejects the null hypothesis when the alternative hypothesis is true, and it’s inversely related to the probability of making a Type 2 error. To test Hitman’s power, we look at the cases where both of *θ*_2_ and *β*_1_ are positive. Here, Hitman’s power is greater than the other methods.

